# Bathymetric and geographical trends in growth of golden redfish (*Sebastes norvegicus*)

**DOI:** 10.1101/739060

**Authors:** Natalia Llopis Monferrer, Benjamin Planque

## Abstract

Golden redfish (*Sebastes norvegicus*) are a long-living (>50 years), late maturing (>10 years) species of commercial interest, distributed along the coast, shelves and continental slope of Norway, down to approximately 400 m depth in the water column. In recent years, analyses of size-at-age data have revealed variable growth trajectories for this species. Whilst some individuals appear to grow slowly after sexual maturity (37cm at ~ 15 years), others continue to grow throughout their lifetime up to 100 cm or more. To investigate how depth and latitude affect golden redfish growth patterns, we developed non-linear mixed effects statistical models. Alongside this, small scale experiments were also conducted to assess the quality of age-determination. The results showed that individuals found in deeper, northern waters present a higher growth potential, even when uncertainty in age determination and species identification were considered. The proximal causes for variations in the growth potential of *S. norvegicus* are still unresolved and the existence of a possible cryptic species remains a fundamental issue that will need to be addressed, in order to understand the causes behind observed growth variations.

## Introduction

Environmental perturbations affect fish growth, with body size varying over geographical and depth gradients (Conover and Present, 1990; Fujita et al., 1995; Robertson et al., 2005; Trip et al., 2014). This has important implications for natural mortality, biomass and catch of individual species, which in turn determine the structure of fish communities (Rountrey et al., 2014).

Within species, larger individuals have been repeatedly observed in deeper waters (Richards, 1986; Gordon and Duncan 1987). Macpherson and Duarte (1991) examined demersal fish communities both in NW Mediterranean and SE Atlantic species, confirming the trend of increased fish size at greater depths for both demersal communities. More particularly, in redfish, it has widely been described that fish migrate to deeper waters as they grow (Atkinson, 1986). These migrations seem to be related with maturation, however, since maturation is correlated with size, bigger fish will be found in deeper waters (Saborido-Rey, 1994). Moreover, nursery areas, where immature fish inhabit, are generally found in shallower waters which will explain the higher abundance of smaller fish in these areas. With regard to geographic distribution, most previous discussions of variation in body size with latitude have focus on Bergmann’s rule. This law states that geographic races of a species found in warmer climates are smaller-sized than those found in cooler areas (Mayr, 1942). Despite the fact this rule is empirical and implies no physiological theory, some studies sustain the temperature as a driving factor (Scholander, 1956). Variations in temperature and photoperiod tend to be more important at higher latitudes (Archibald et al., 1981). Cooler temperatures may allow greater efficacies in energy transfer rates between trophic levels. Furthermore, northernmost individuals tend to be larger to compensate for the shorter growing season, with higher growth rates (Love *et al.* 2002; Carmona-Catot *et al.* 2014). However, this trend runs contrary to some published studies where bigger individuals are found southward (Yamamoto et al., 2010), which is not necessarily translated into overall smaller growth rates (Carmona-Catot et al., 2014). Low-latitude regions provide preferable conditions for processing and assimilating food (Trip et al., 2014) while food supplies are generally less abundant in cool regions (Lindsey, 1966).

Many hypotheses have been proposed to describe variations in fish growth and size. Since most fish species are poikilotherms (i.e. they do not regulate their body temperature), ambient temperature can influence a number of individual traits, including metabolic rates and growth rate as well as spatial and vertical distribution (Allan and Castillo, 2007). Physiological constraints due to temperature can, directly or indirectly, drive latitudinal and depth gradients in fish body size and growth but the generality and mechanisms underlying these trends are unclear and controversial.

Fish growth indicates the energy production within the individual and it arises from the allocation of energy between an increase in size (length) and an increase in condition (i.e., gram per year) (Webber and Thorson, 2016)). This parameter is sensitive to environmental perturbations, it is primarily determined by the amount of energy acquired by individuals (Longhurst, 2010). The amount of energy allocated to growth will depend on a number of factors and will influence all levels of biological organisation. Some of the factors are intrinsic (genetically and physiologically) while others are driven by environmental factors such as temperature, oxygenation levels and feeding (Dutta, 1994; Boeuf and Payan, 2001; Saborido-Rey and Kjesbu, 2005; Magnussen, 2007; Enberg *et al*., 2008). Energy acquisition largely depends on access to prey and their nutritional quality. Resource availability varies in time and space and can lead to variable growth, rapid growth occurring when and where food availability is high, while reduced or even negative growth occurs when and where food is scarce. During early life stages, energy is needed for development and rapid growth. At maturity and with increasing body sizes, individuals invest decreasing proportions of their increasing total energy intake to growth, energy is now rather channelled towards reproduction and gonad development, resulting in a lower metabolic rate (Paloheimo and Dickie, 1966). Changes in body size across depth and temperature gradients reflect a combination of changes in age composition and in growth rates and can therefore be difficult to interpret. Variations in body size-at-age directly reflects changes in somatic growth patterns.

The *Sebastes* genus comprises approximately 110 species worldwide (Hyde and Vetter, 2007) and most *Sebastes* species are found in the Pacific Ocean, where they exhibit great morphological variability. In comparison, only four species of this genus inhabit the North Atlantic Ocean. Within the genus *Sebastes*, the phenomenon of increasing fish size with increasing depth has been commonly observed (Lenarz, 1980; Wilkins, 1980). In Pacific waters, Richards (1986) documented an increase in size with depth for three species of rockfishes (*S. elongatus*, *S. maliger* and *S. ruberrimus*). Ingram (2011) however, analysing individuals from 66 rockfish species, did not observe any significant correlations between body length and depth.

Members of the *Sebastes* genus are known to be generally slow-growing (Sandeman, 1961) and long-lived (Archibald et al., 1981). Their somatic growth is variable within the genus, with some species presenting determinate growth (*S. mentella*) while others have indeterminate growth and continue to grow after maturation, throughout their lifetime (*S. norvegicus*).

This study focuses on the golden redfish (*S. norvegicus*) in northeast Atlantic Ocean. Golden redfish is long-lived (up to 65 years; Campana et al., 1990) and slow growing (max length, 100 cm). The analysis of length-at-age data has revealed variable growth trajectories (Figure 1). Juvenile stages seem to follow a common, almost linear growth pattern. After maturity (around 11 years) length-at-age appear to vary greatly between individuals. While some individuals display reduced growth, with adult size remaining around 40 cm, others can reach sizes over 60 cm at ages beyond 25 years. It is yet unclear if these different patterns in length-at-age are related to geographical/latitutdinal and depth gradients. To examine how golden redfish observed growth rates vary in response to latitudinal and depth gradients, we used empirical observations from fisheries and research surveys conducted in the Norwegian and Barents Seas over the last two decades. We then modelled growth using hierarchical models as an approach to determine whether there is a general tendency towards increasing size-at-age in deeper waters and higher latitudes.

**Figure 1:**
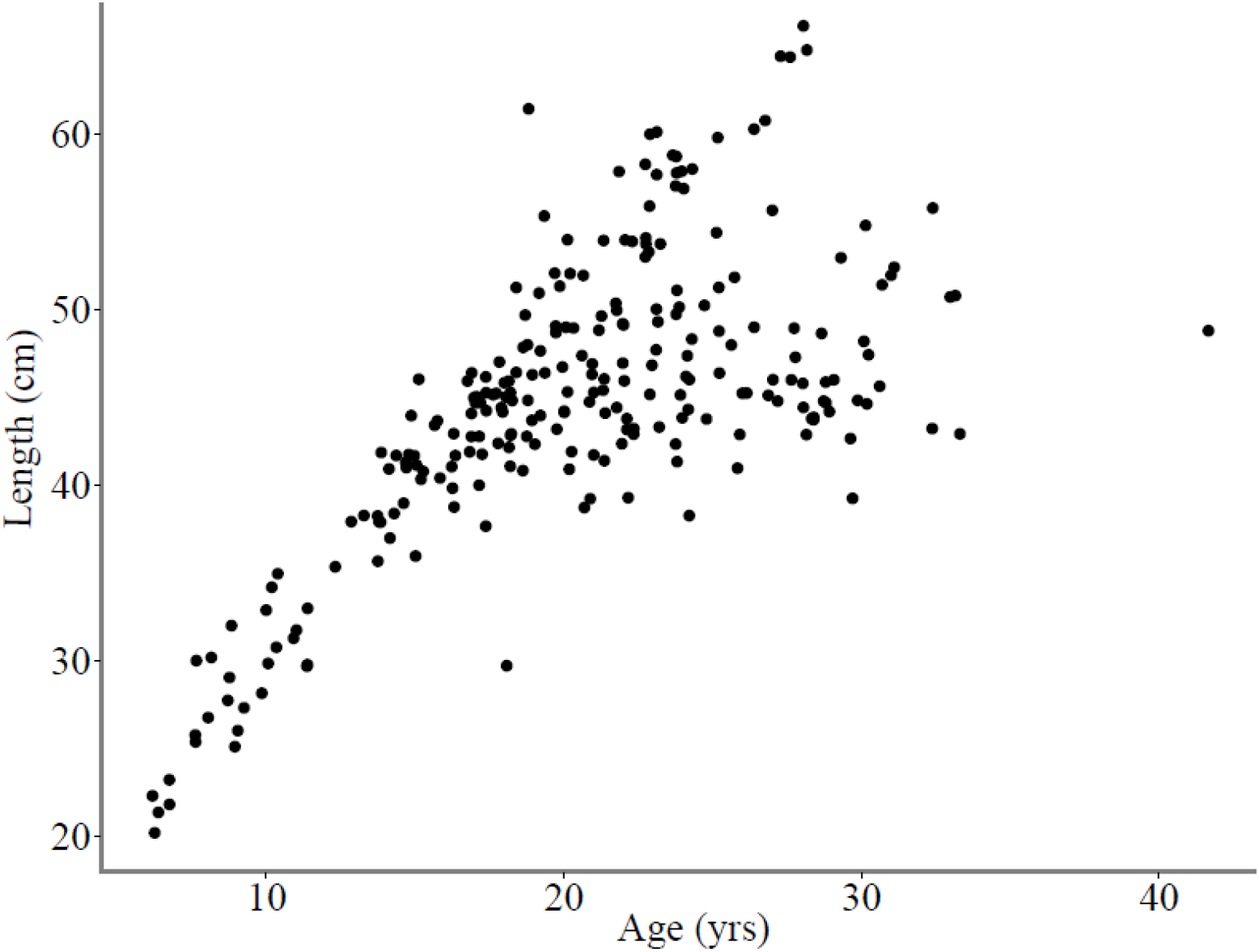
Length-at-age of 248 individual specimens of *S. norvegicus* collected in 2013. After maturation (11 years old), some individuals show reduced growth reaching a maximum length of approximately 40 cm while other individuals can reach lengths over 60 cm.

## Material and methods

### Biological data

Age and length data from 13056 fish (6322 males and 6734 females) identified as *S. norvegicus* were extracted from the database held by the Institute of Marine Research of Norway. Samples were collected with bottom, shrimp and pelagic trawls, gillnets and longlines by research and commercial vessels in the Norwegian and Barents Seas between 1992 and 2015. Specimens were obtained following the IMR manual for sampling fish and crustaceans (Mjanger et al., 2016). Data for each individual included geographical coordinates, date and time of sampling, fishing gear used, depth and duration of sampling, fish length, age, sex and maturity stage. For bottom trawls, sampling depth was set to ocean floor depth at the sampling location. For pelagic nets and trawls, sampling depth was defined as the mean of the gear deployment depth recorded during fishing. Species identification was performed on-board based on body size, beak size, eye diameter and direction of spines in the pre-operculum, as suggested by Barsukov *et al*. (1984). For few individuals, morphological identification was supplemented by lab-based genetic species identification. Total fish length was measured on board and reported to the centimetre below. Maturity stage was determined according to the maturity scale in Mjanger *et al.* (2016). Age determination was conducted using the otolith break-and-burn technique by counting the hyaline zones under the microscope (winter zones; dark in reflected light, ICES, 2009b).

There was no direct registration of environmental parameters (e.g. temperature, prey availability, predatory pressure or competition) simultaneous to fish sampling. To investigate the potential effects of environmental conditions on growth, we used depth and geographical location as proxies of the environmental conditions experienced by individual fish.

### Growth models

Phenomenological (and statistical) growth models, describe growth as a function relating individual length to individual age (and possibly other factors). The most studied and commonly applied statistical models are: the von Bertalanffy growth function (VBGF), the Gompertz growth function, the logistic model and Schnute-Richards functions (Katsanevakis, 2006). The VBGF has been widely used to describe indeterminate fish growth (e.g., Czarnoleski and Kozlowski, 1998). Paradoxically, although the biological rationale behind the equation has been shown to be erroneous, the VBGF fits well to many observations both for individual growth trajectories and for population averages (Enberg et al., 2008). The VBGF model tends to work well for adult individuals in which surplus energy is channelled towards reproduction but less so for juveniles (Lester et al., 2004). We modelled growth using the VBGF which expresses the length as a function of age:

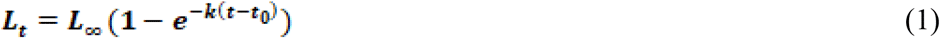

where *L_t_* is the expected or average length at time *t*, *t*_0_ is used to adjust the model for the initial size of the animal, *k* represents the exponential rate of approach to the asymptotic size, and *L_∞_* is the asymptotic maximum length. In this formulation of the VBGF, the parameters *L_∞_* and *k* are strongly correlated and might therefore be difficult to estimate. Alternative parametrisations of the VBGF have been proposed, in which parameters are less correlated or at least that are better suited for parameter estimation by computer optimization. For example, Gallucci and Quinn II (1979), introduced a new parameter, ω = *kL_∞_* which, when substituted in equation 1, results in a new parametrization:

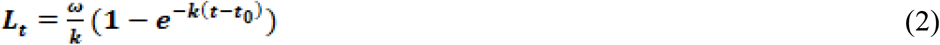

The Schnute parametrization (Quinn II and Deriso, 1999) is based on “expected values” at known ages and is also intended to lower the correlation between parameters:

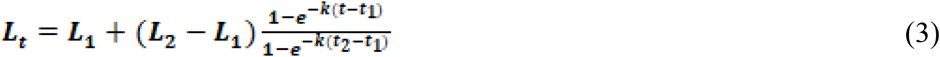

where *L_1_* is the average length at the youngest age (6 years for both males and females) in the sample, *t_1_*, and *L_2_* is the average length at the oldest age (45 years for males and 54 years for females) in the sample, *t_2_*. In the Schnute parametrization, the estimated parameters (*L_1_, L_2_, t_1_* and *t_2_*) are more directly interpretable than in the original formulation (equation 1) in which *t_0_* and *L_∞_* are not observable quantities. In the present study, we used the Schnute parametrization (equation 3) which proved efficient for parameter estimation. We found that parameters were not less correlated than with the other parametrizations.

### Effects of latitude and depth on growth

To explore which external factors could be affecting growth, several statistical models were built considering the following variables: depth and region. We built hierarchical models where the growth hyper-parameters (*L_2_*, *L_2_* and *k*) were common to the whole population and in which variations in growth were specific to particular groups of individuals. We used nonlinear mixed effect-modelling (NLME, Pinheiro and Bates, 2000), where fixed effects represent what is common to the population while random effects are specified for each group. Given that the VBGF is best suited to represent the growth of mature individuals, only data from mature fish were selected to model growth. Models for males and females were constructed independently to account for the possibility of sexual dimorphism.

Sampling depths were divided into seven intervals (Table 1) and geographical locations were divided into eight polygons that covered the study area (Figure 2). Depth intervals and geographical polygons were selected to ensure relative balance in sampling intensity in each category and minimum redundancy between depths and regions (i.e. each region covers several depth intervals and vice-versa).

**Table 1:** Depth intervals used to develop the models.

| Depth interval | Min depth (m) | Max depth (m) | Number of individuals |
| --- | --- | --- | --- |
| 1 | 0 | 170 | 1805 |
| 2 | 170 | 200 | 1919 |
| 3 | 200 | 235 | 2116 |
| 4 | 235 | 270 | 1976 |
| 5 | 270 | 300 | 1894 |
| 6 | 300 | 375 | 2052 |
| 7 | 375 | - | 1294 |

**Figure 2:**
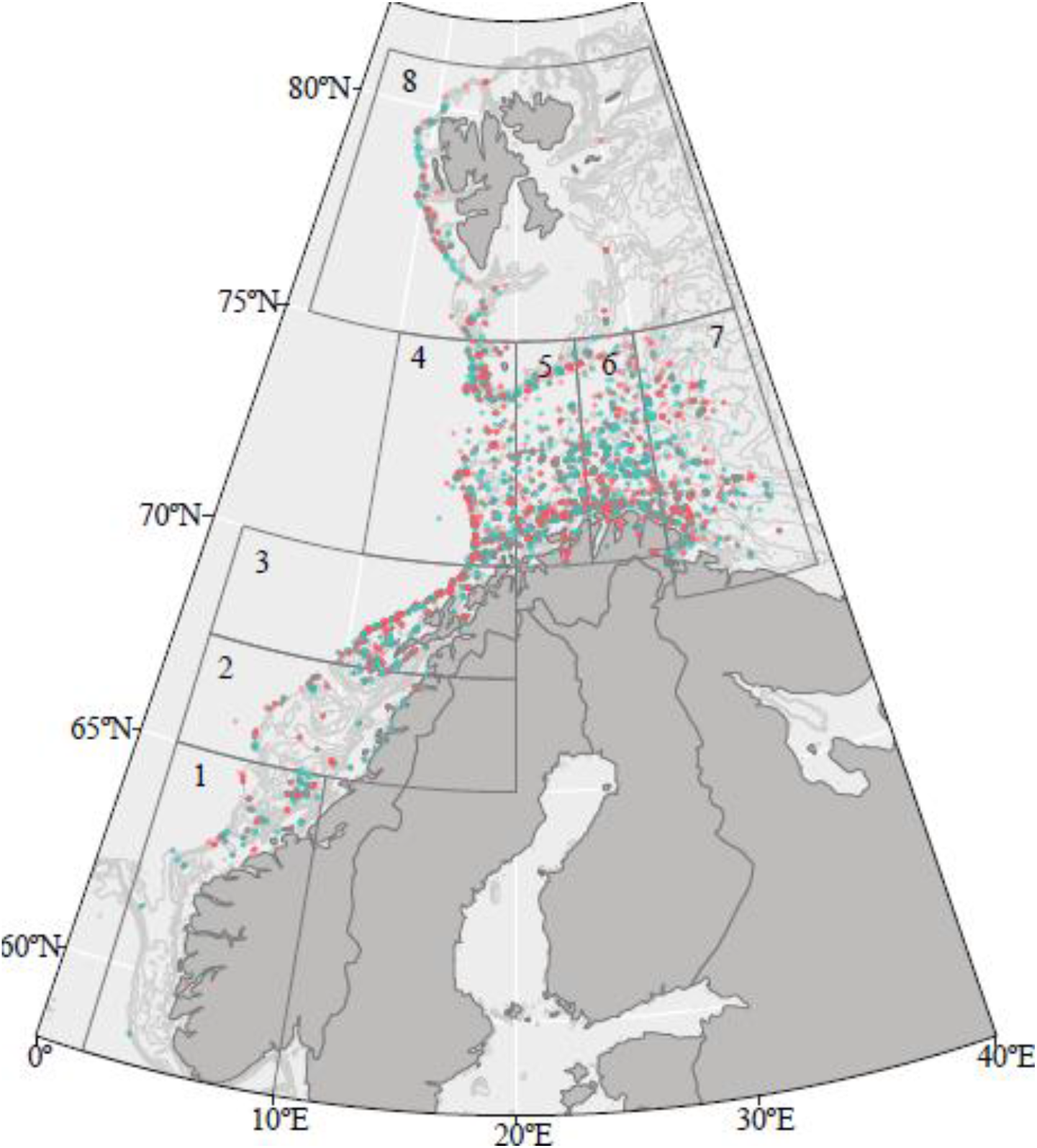
Map of the study area. Grey lines show the isobaths that delineate the depth intervals used in the hierarchical models. Polygon numbered from 1 to 8 delineate the geographical regions used in the models. Observations of male and female fish are shown in blue and orange respectively.

We constructed single-level and multi-level models to study the effect of depth and regionalisation. Single-level models consisted in the study of the individual effect of one of the parameters (e.g., equation 4: length ~ age | depth: parameters of the VBGF vary between depths intervals), while the latter, analysed the hierarchical combination of both parameters making it possible to establish a prioritization of them (e.g., equation 5: length ~ age | depth/region: parameters of the VBGF vary between depths intervals and within each depth interval they also vary between regions).

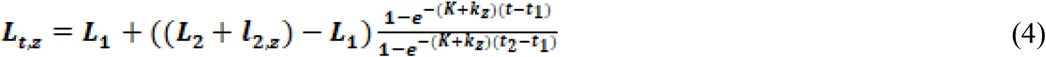

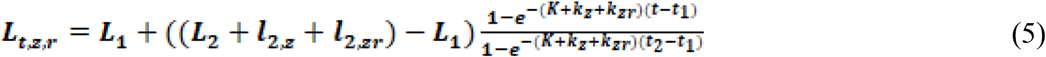

*L_1_*, *L_2_* and *K* are the hyper-parameters of the model. *l_2,z_* and *k_z_* are the random effects for a given depth interval in the single-level model and and are the random effects for a given region within a specified depth interval.

Model parameters were estimated using the nlme library (Pinheiro et al., 2017) in R (R Core Team, 2016).

The percentage of variability explained by the model was calculated using equation 6.

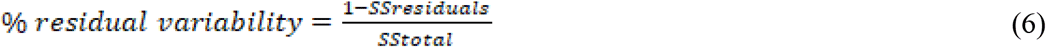

where *SStotal* is given by 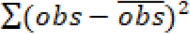 and *SSresiduals* is given by 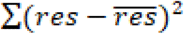, *res* being the difference between observations and predictions.

## Results

### Growth modelling

Since sexual dimorphism could be a cause of variations in observed growth, we modelled growth of males and females separately. Results showed that there exist slight differences between male and female size-at-age. Length at maximum age was generally higher for females than for males (Table 2, Figure 3A and 4A). However, these differences were usually less pronounced that those attributable to depth or region and in few instances, *L_2_* was smaller for females than for males.

**Table 2:**
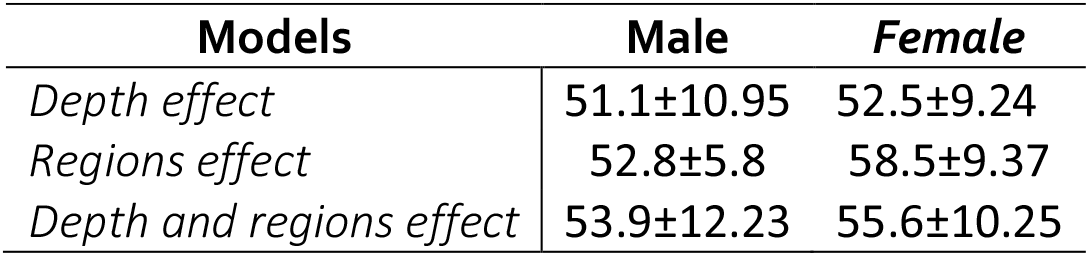
*L_2_* values and standard deviation for each model and sex.

**Figure 3:**
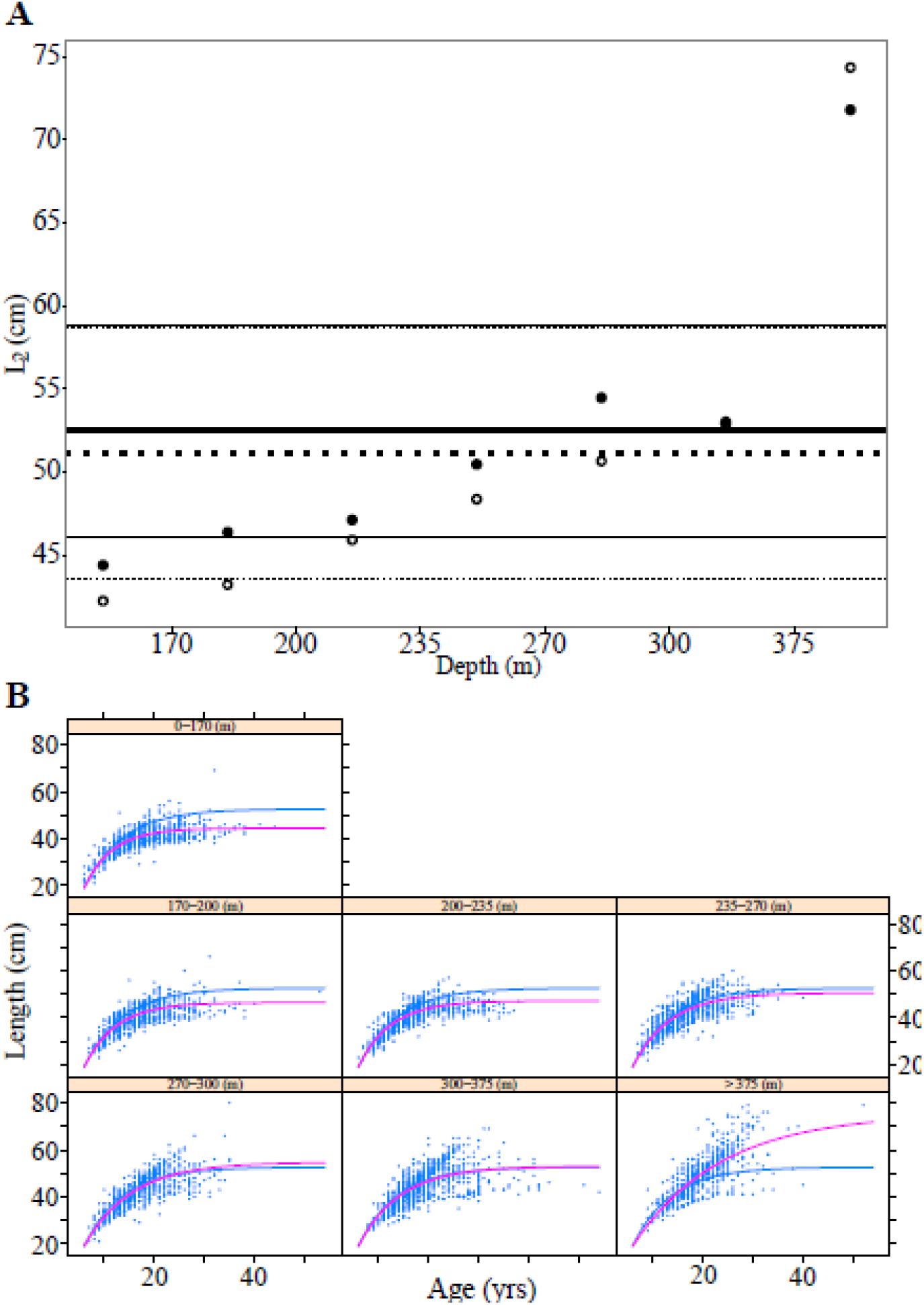
VBGF models with depth effect. A. Horizontal lines (solid for females and dotted for males) indicate the mean (thick lines) and 95% confidence intervals (thin lines) of *L_2_* for the whole population (fixed effects only). Dots (filled for females, empty for males) show *L_2_* + *l_2,z_* values estimated for each depth interval (fixed and random effects). B. Fitted VBGF model considering the effect of depth for females. Panels are for the different sampling depths. Blue dots: observations, blue lines: model with fixed effects only, pink lines: mixed model with depth effect.

### Model with depth effect

In the single-level model where the VBGF parameters vary between depth intervals, mean length at maximum age for the females/males is estimated to be 53/51 cm. Individuals show a pattern of increasing length with increasing depths, to the exception of the depth interval of 300-375 m where a slight decrease can be observed. In the last depth interval there is an important increase of more than 20 cm (Figure 3A). The data displayed in Figure 3B are the results of statistical growth modelling for female individuals. Within each depth interval, the variability not explained by the model amounts to 33%. The hyper-parameters *L* and *k* are negatively correlated. The model with depth effects shows definitely an increase with depth, with an important increase in the deepest layer. Model results for the male fraction of the population are presented in the Supplementary material.

### Model with regions effect

In the single-level model where growth varies between regions, mean length at maximum age for females/males is 58/54 cm. In northern regions, values are higher than in southern regions (Figure 4A). The residual variability amounts to 39%. Figure 4B shows the results of statistical growth modelling for female individuals with latitude changes. Fast and slow growing individuals are found at each region except in region 8, situated in the northern area, where only fast growing individuals are found. Regions 4, 5, 6 and 7 are located in the same latitude. While regions 4 and 5 present values of length at maximum age (*L_2_*) lower than the mean (fixed effects), values of this parameter found for regions 6 and 7 exceed the mean (Figure 4B). Model results for the male fraction of the population are presented in the Supplementary material.

**Figure 4:**
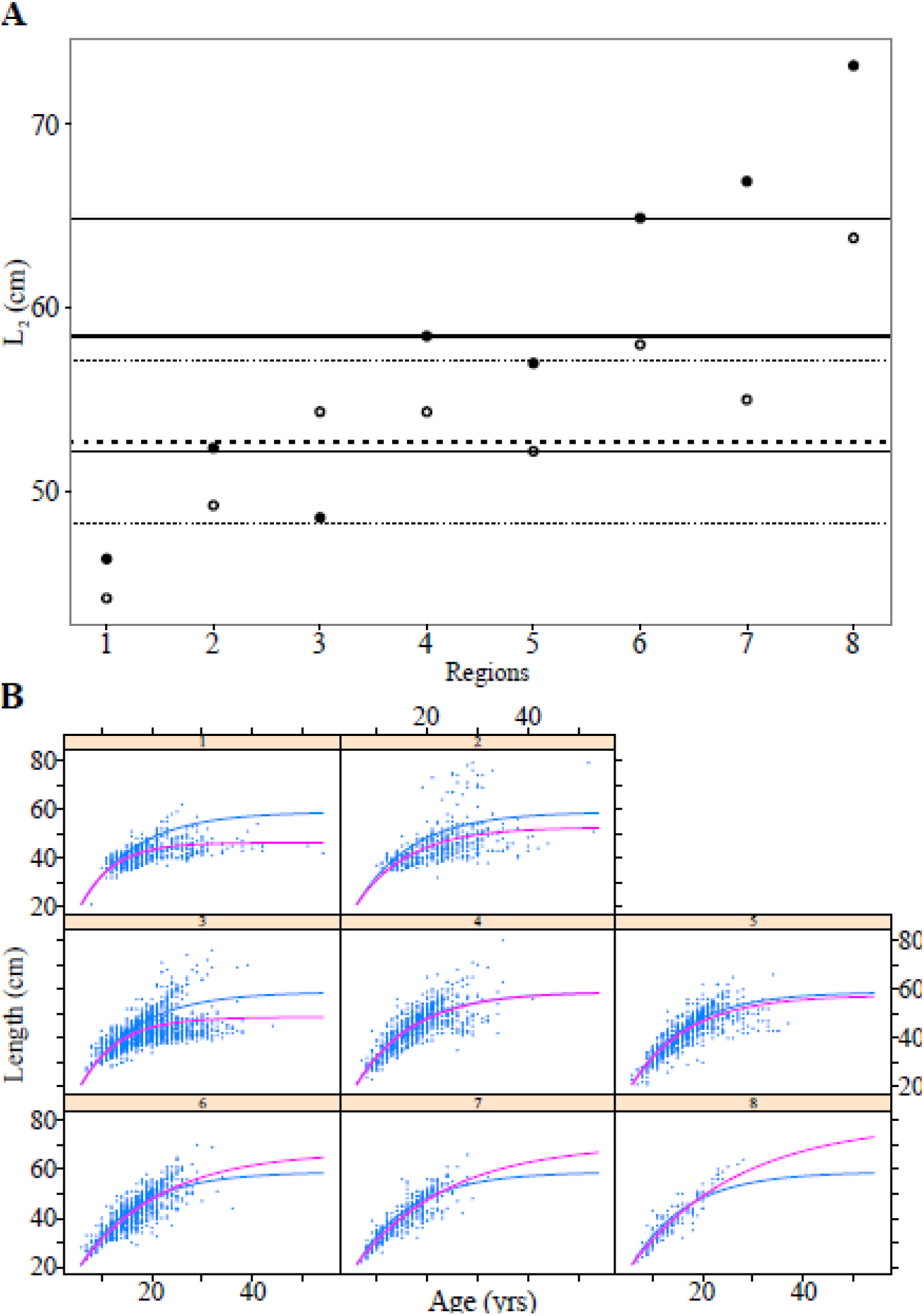
VBGF models with regions effect. A. Horizontal lines (solid for females and dotted for males) indicate the mean (thick lines) and 95% confidence intervals (thin lines) of *L_2_* for the whole population (fixed effects only). Dots (filled for females, empty for males) show the *L_2_* + *l_2,z_* values estimated for each region (fixed and random effects). B. Fitted VBGF model considering the effect of depth for females. Panels are for the different regions. Blue dots: observations, blue lines: model with fixed effects only, pink lines: mixed model with depth effect.

### Hierarchical model with depth and regions effect

The hierarchical model combines depth and regions as sources of random variation. In this case, depth is placed at the first level (model results with region at first level are presented in the supplementary material). The length at maximum age estimated through this multi-level model was of 56/54 cm. When random effects are considered, the greatest lengths at maximum age are found in the deepest and northernmost regions (Figure 5A), confirming the results from the single-level models. The residual variability found in this model is slightly lower (28%) than the one found for the single-level models, both for depth and regions (Figure 5B). Male results can be found in the Supplementary material.

**Figure 5:**
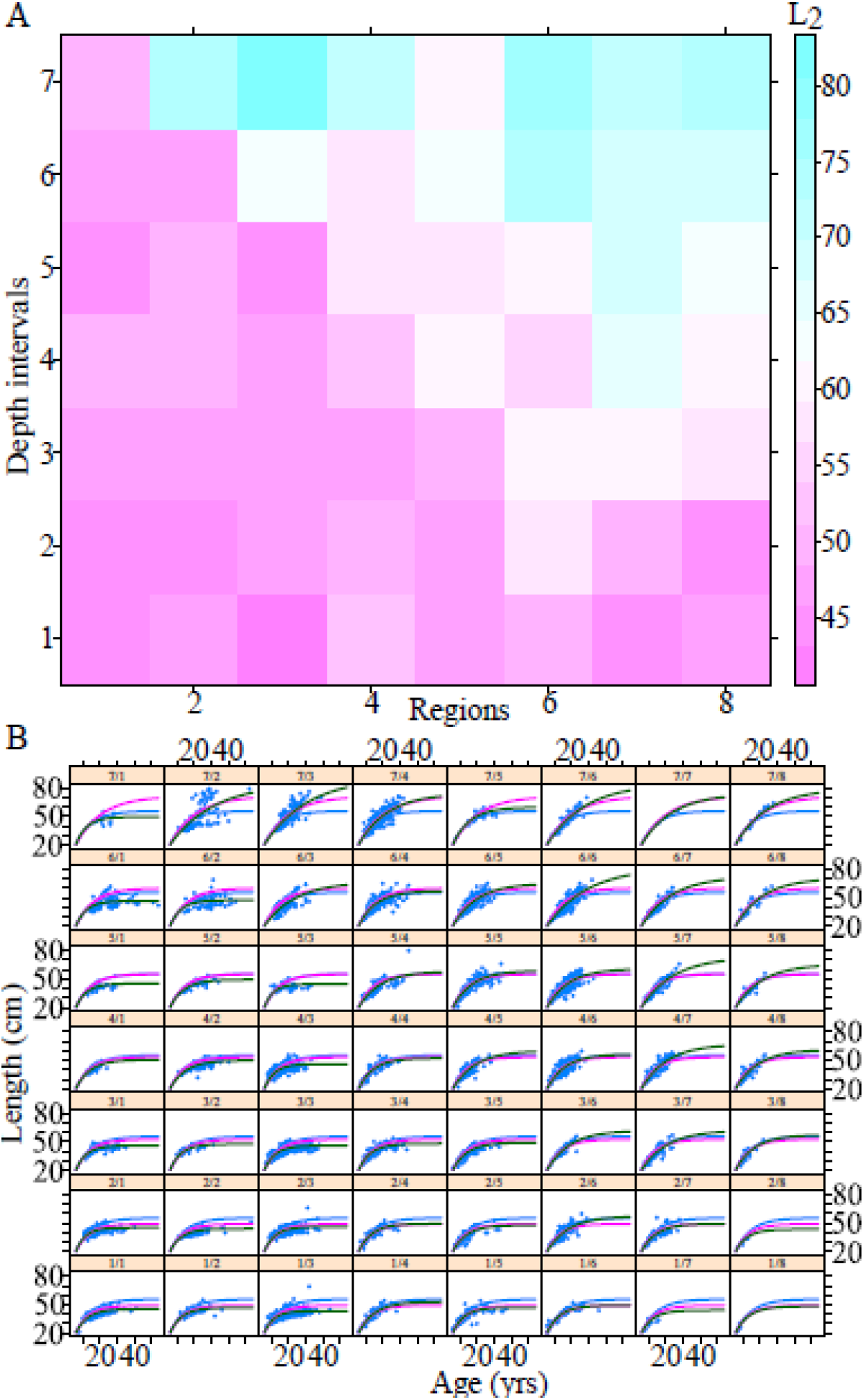
VBGF models with depth and region effect for females. A. Matrix of *L_2_* values estimated for *S. norvegicus* for each combination of depth intervals and regions. B. Fitted VBGF model considering the multi-level effect. Panels are for each combination depth/regions. Blue dots: observations, blue lines: model with fixed effects only, pink lines: mixed model effect with depth, green lines: hierarchical mixed models with depth at first level and regions at second level.

## Discussion

In this study, we have explored different, and non-exclusive, hypotheses that can explain observed variations in the growth of *S. norvegicus.* These include sexual dimorphism, taxonomic misidentification, bias and uncertainty in age determination and variations in environmental conditions. While sexual dimorphism exists in *Sebastes*, differences in size-at-age between males and females are too small to explain observed growth patterns. On average *L_2_* values estimated for females were only slightly greater than the *L_2_* values found for males in all models, in accordance with published results (Gascon, 2003; Cadigan and Campana, 2016), but we also found that males had a higher growth potential than females at greater depths.

For this study, species were identified on-board using morphological criteria only, knowingly that morphological species identification within the genus *Sebastes* is difficult (Magnússon, 1981). *S. mentella* is generally found in deeper waters than *S. norvegicus* (Stransky, 2005) and *S. mentella* grows smaller than *S. norvegicus* (Garabana, 2005). If species misidentification was systematic and a large fraction of the specimen studied were *S. mentella* (but wrongly identified as *S. norvegicus*), we would expect to observe a large fraction of smaller fish in deeper waters, which is in opposition to the results of this study. This indicates that the results of the present study are robust to potential errors in taxonomic identification.

The statistical approach used here makes the implicit assumption that there was no bias in age determination. It is known, however, that age determination can be challenging in *Sebastes* species (Campana, 2001; Johansen, 2003). Results from the double-age reading experiment showed that there can be substantial variations between multiple determinations of age of *S. norvegicus* conducted by the same otolith reader. The uncertainty estimated in age amounts to 3.8%. The age determination conducted in 2017 resulted in more variable ages than those determined in 2013. Because age reading in 2017 was made using the same part of the otolith that was broken and burnt in 2013, possible erosion of the hyaline zones over time may have led to a harder and less precise reading. This can partially explain the differences between the first and second age readings. There was no systematic bias in age readings between the first and second readings. Given these results, it seems unlikely that individuals identified as slow growing may primarily result from systematic misreading of age, i.e. with overestimated age that would result in under-estimated growth. Rather, regional and bathymetrical variations in growth appeared despite relatively high uncertainty in age determination. This result suggests that these variations are real and that they would have been better identified, had uncertainties in age determination been reduced.

This investigation revealed a higher growth potential for *S. norvegicus* individuals from deeper waters than those from shallower waters. Likewise, *S. norvegicus* from northern latitudes presented also a higher growth potential compared to individuals found in southern latitudes. The effect of water depth on growth of *S. norvegicus* is little documented but it has been reported that smaller fish from the genus *Sebastes*, tend to live in shallower waters (Lenarz, 1980; Wilkins, 1980), which is consistent with current results. Individuals observed in northern latitudes were not the largest on average, since they were generally younger than individuals from southern latitudes. However, they showed a higher growth potential than fish located in southern regions. Conover and Present (1990) suggested that growth and other life history traits in fishes can vary between latitudes. The driving selective force for this variation in growth was demonstrated to be size-dependent winter mortality in young-of-the-year fish (age-0 fish), which is higher in smaller fish (Shuter and Post, 1990) and at higher latitudes (Conover and Present, 1990). The high mortality in small individuals due extreme environmental conditions in northern latitudes may result in a strong selection pressure towards fast growing fish with increasing latitude.

Depth and regional variations may constitute proxies for other abiotic and biotic factors such as salinity, temperature, food availability or predatory pressure which all can contribute to changes in fish growth. Consequently, although results from the non-linear mixed-models indicate that growth potential varies with depth and geographical regions it is not possible to conclude that these external factors (depth and regionalization) are the proximal causes for variations in growth or that these are the only factors responsible for variations in growth.

Even though this study has revealed that observed variations in growth potential can partially be explained by depth and geography, it cannot be specified whether these variations result from environmental effects on fish growth or from spatial and bathymetric variations in the relative distribution of the different genetic types of *norvegicus*. Two distinct cryptic species, *S. norvegicus*-A and *S. norvegicus*-B, have been claimed by Saha *et al*. (2017) and observed variations in growth patterns may possibly reflect the existence of genetically separated subpopulations that display different growth rates and are preferentially located in different depths and regions. Regular genetic analyses of specimens collected by commercial and research vessels are needed to explore this cryptic-species-hypothesis.

## Conclusion

In summary, we found that differences in growth rates between sexes are minor and cannot explain observed variations in fish growth. We also found that uncertainties and potential biases in age determination and species identification are also unlikely to generate the growth patterns observed. Rather, variations in growth are associated with depth and regions (higher growth rates for northern and deeper fish) but the mechanisms causing these variations remain unresolved. It is possible that these variations are caused by the direct effect of the bathymetry and geographical location although it is more likely that these two parameters are only proxies for other factors such as temperature, salinity, habitat quality, prey availability or predation pressure. Finally, the issue of potential cryptic species (genetically different but morphologically counfounded) having different growth rates and different spatial and bathymetric distributions remains unresolved.

## Acknowledgements

Thanks to the IMR and commercial vessels for the sampling collection. We wish to give our gratitude to Kjell Nedreas and Arne Storaker from Institute of Marine Research (IMR) in Bergen, for their technical work and the rapidity with which they responded to our requests and to Torild Johansen from IMR in Tromsø for her knowledge and work in the field of genetics. Many thanks also to Professor Gilles Yoccoz and Uffe for their constructive comments and useful remarks improving our work.

## Supplementary material

### VBGF models results for males

Plots of Von Bertalanffy Growth functions (VBGF) with depth effect and regions effect. Panels are for the different sampling depth intervals and for the different regions. Blue dots correspond to the observations. Blue line represents the model with only fixed effects and pink line represents the mixed model effect with the depth or region effect.

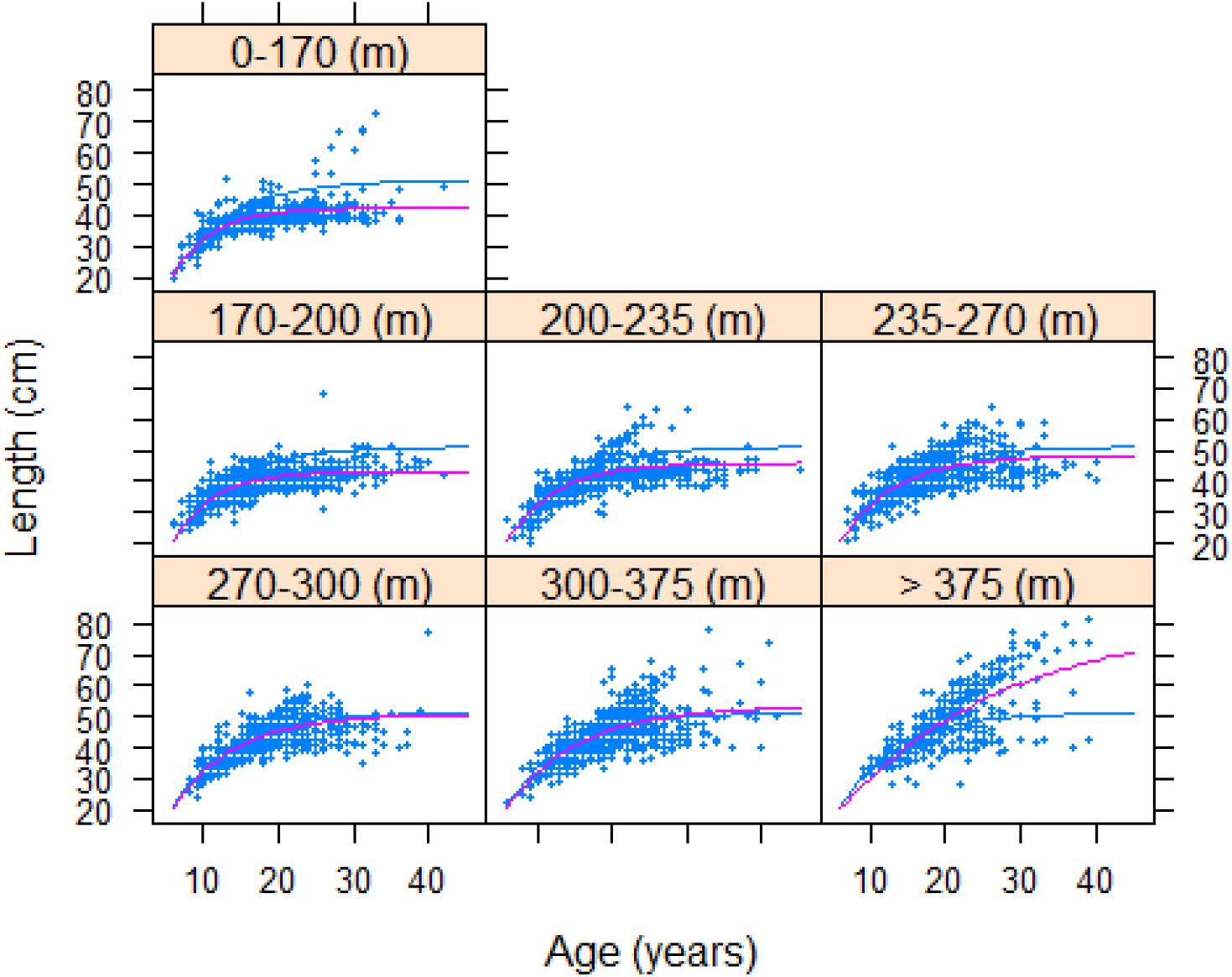

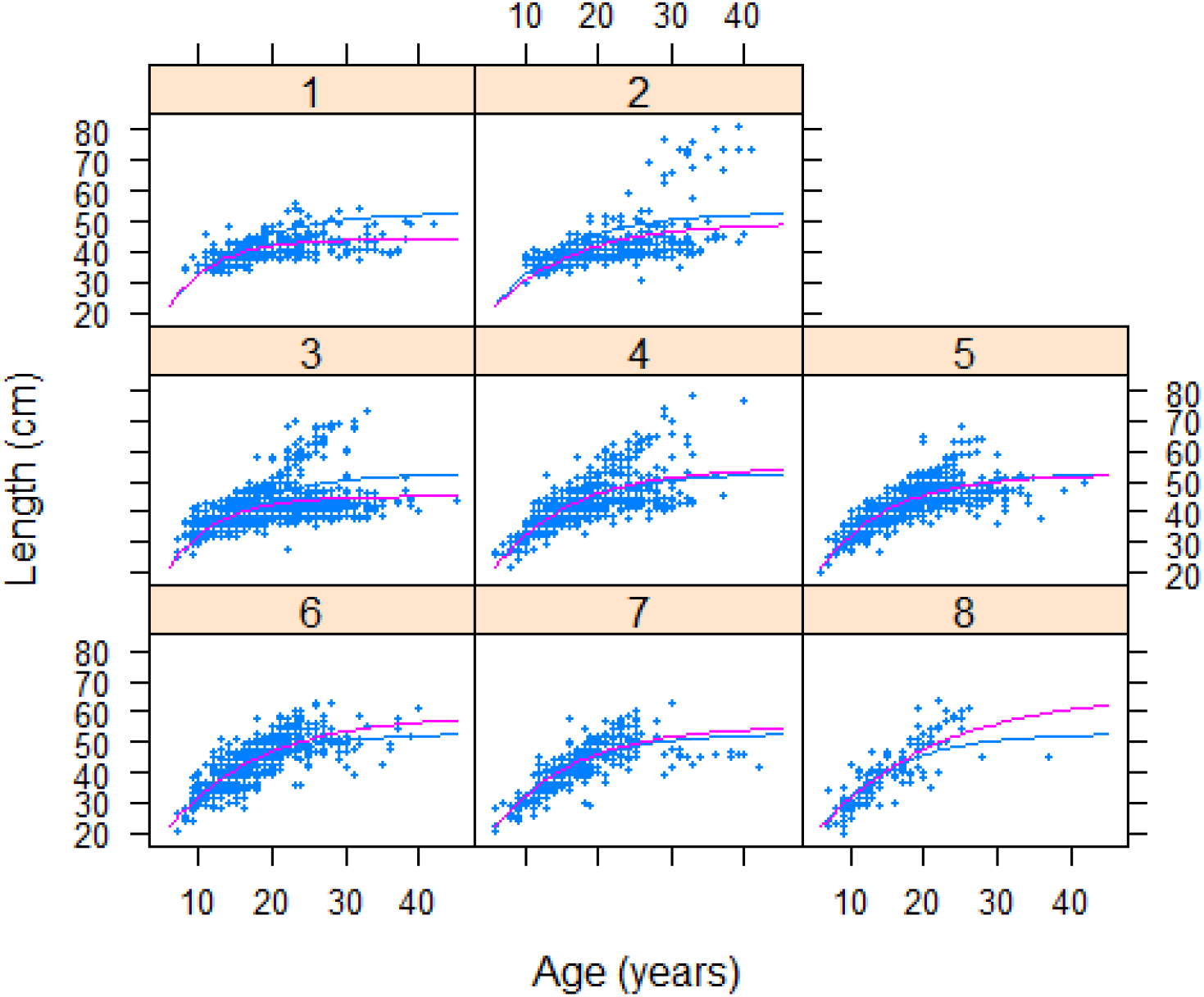

VBGF models with depth as a main factor in the hierarchical model and region as the second factor. Matrix of *L_2_* values estimated for *S. norvegicus* for each combination of intervals of depth and regions.

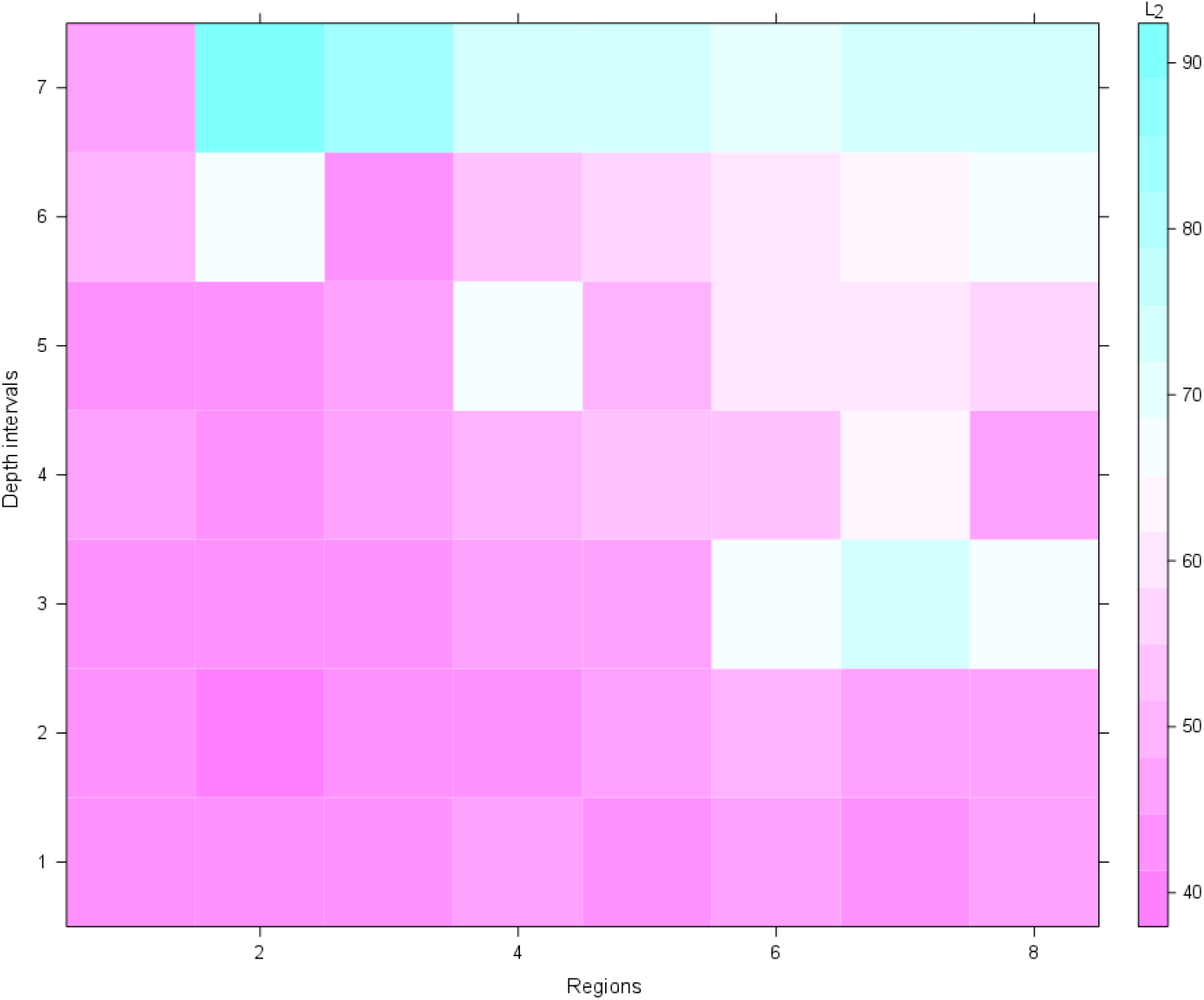

VBGF models with depth as a main factor in the hierarchical model and region as the second factor. Panels are for each combination depth/regions. Blue dots correspond to the observations. Blue line represents the model with only fixed effects common to the whole population. Pink line represents the mixed model effect with the depth (fixed and random effects). Green line represents the hierarchical mixed model effect with depth as the first level and regions as the second one (fixed and random effects).

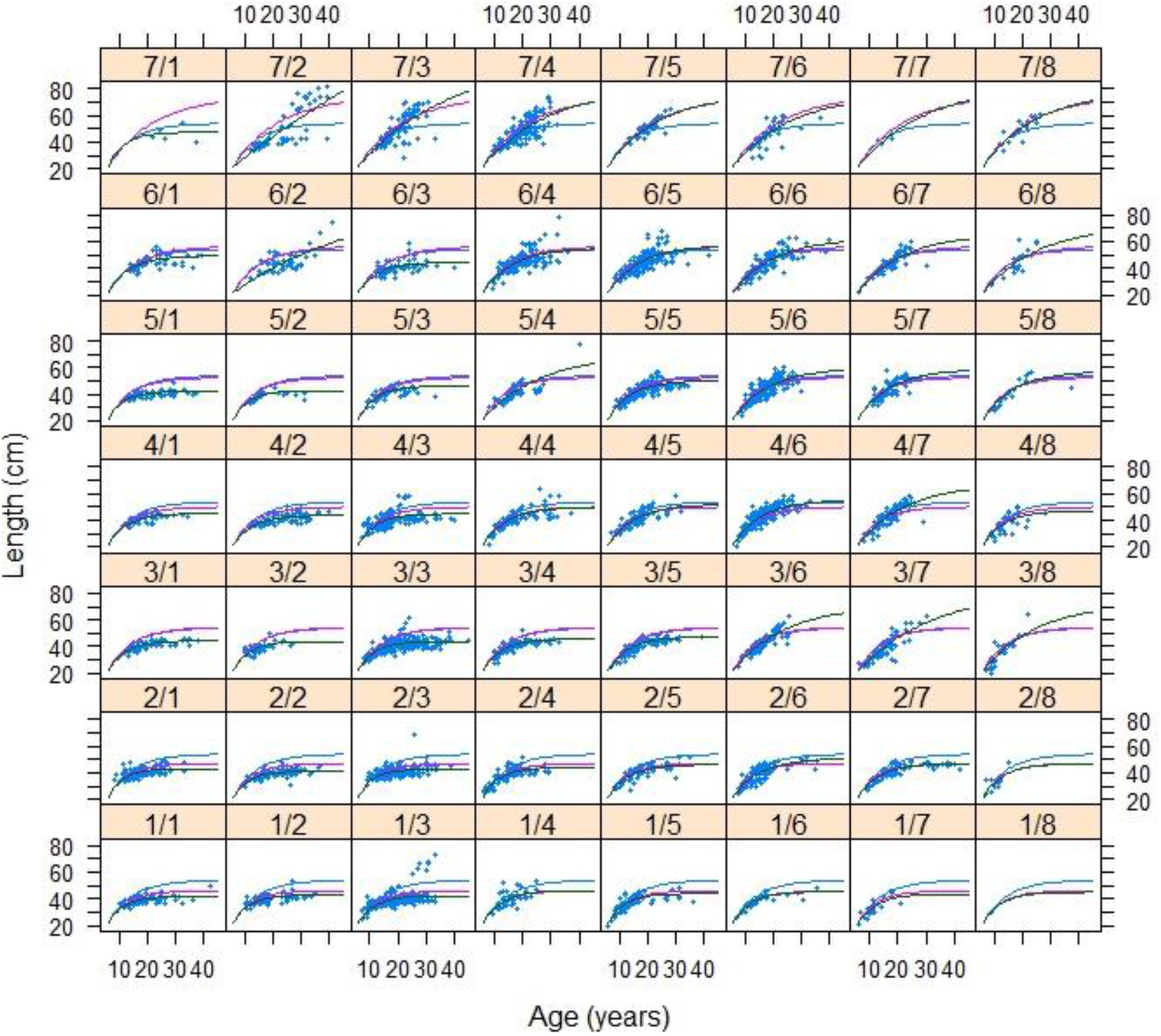

VBGF models with region as a main factor in the hierarchical model and depth as the second factor. Matrix of *L_2_* values estimated for *S. norvegicus* for each combination of intervals of regions and depth.

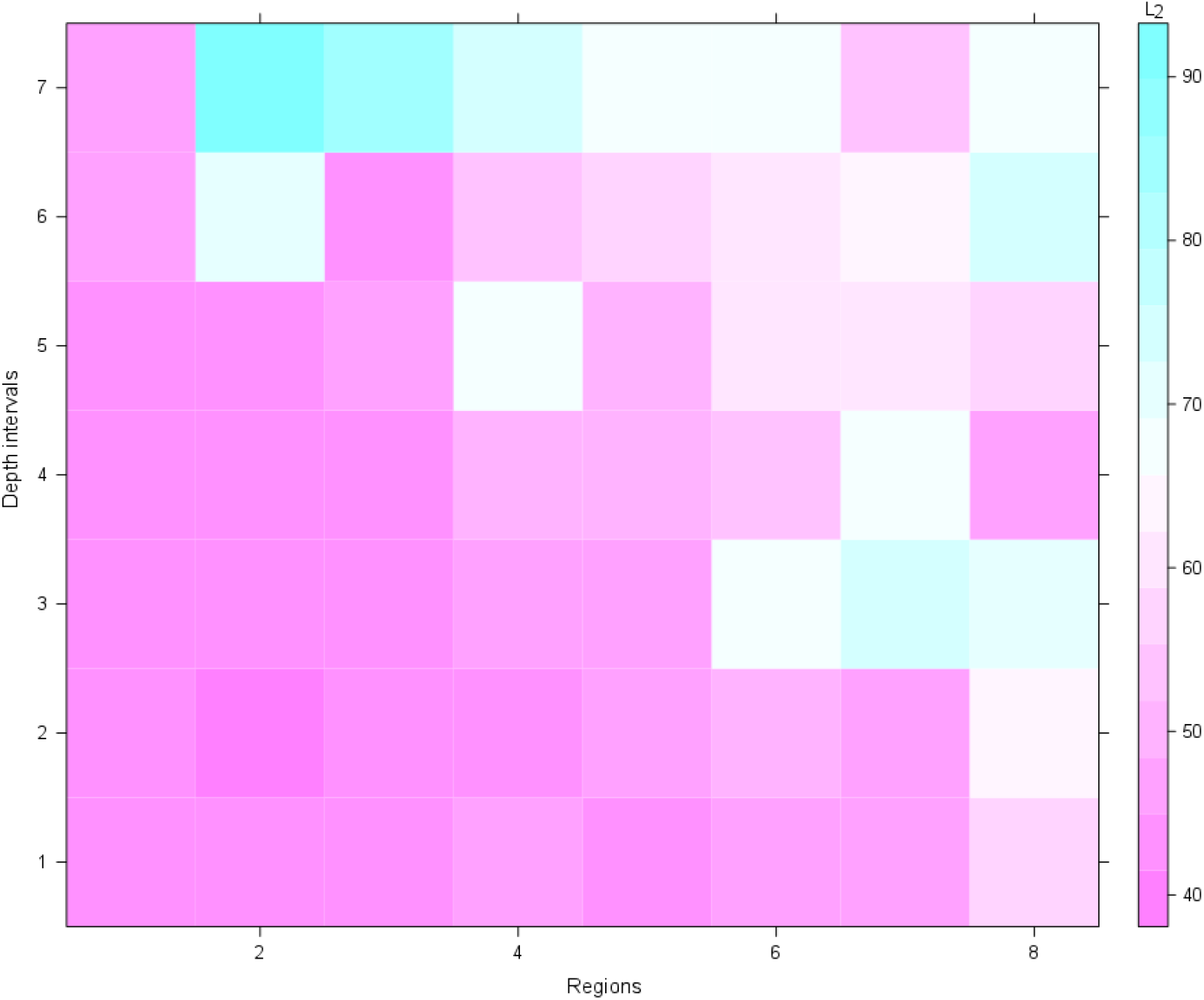

VBGF models with region as a main factor in the hierarchical model and depth as the second factor. Panels are for each combination of intervals of region/depth. Blue dots correspond to the observations. Blue line represents the model with only fixed effects common to the whole population. Pink line represents the mixed model effect with the region (fixed and random effects). Green line represents the hierarchical mixed model effect with regions as the first level and depth as the second one (fixed and random effects).

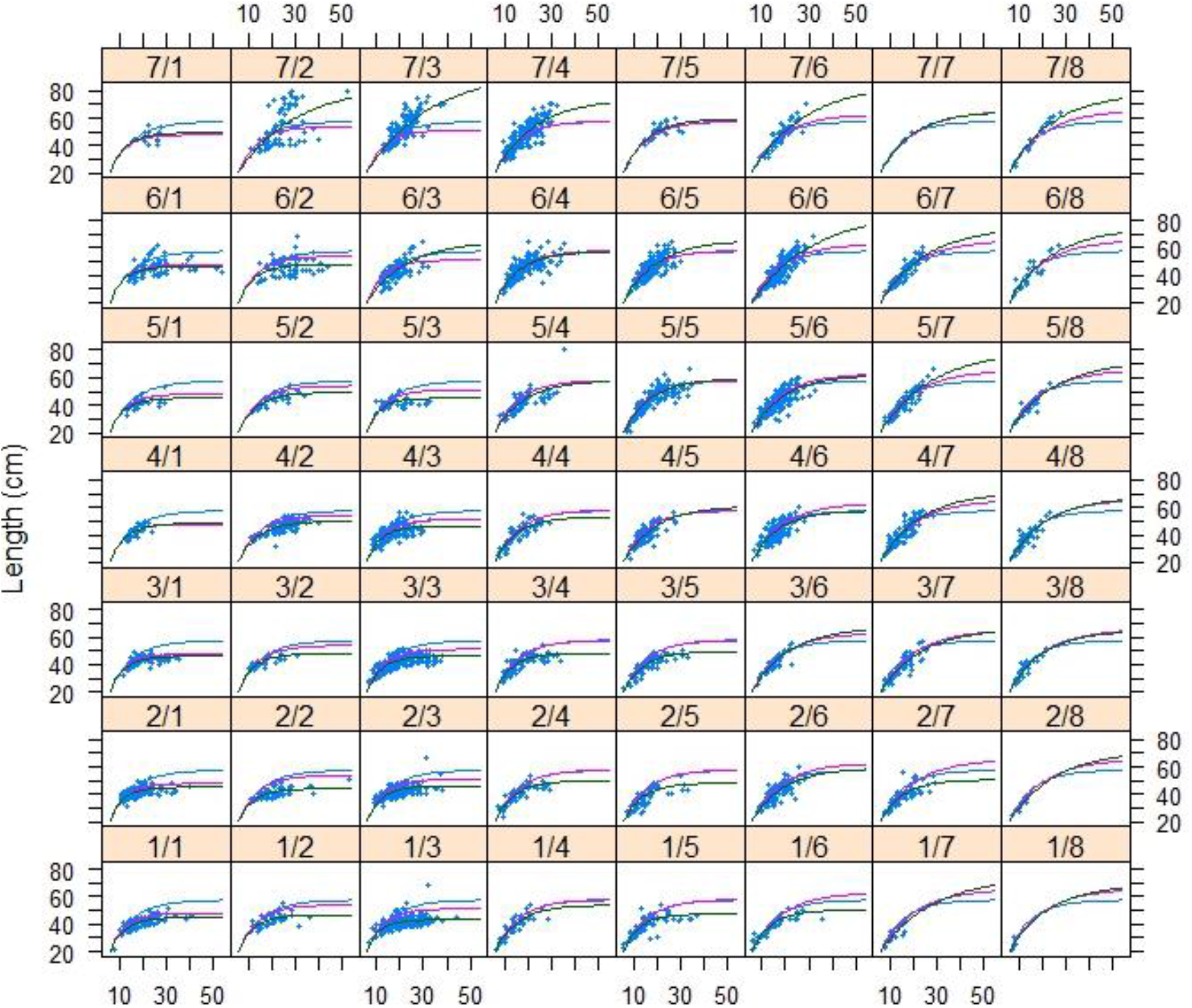

### VBGF models results for females

VBGF models with region as a main factor in the hierarchical model and depth as the second factor. Matrix of *L_2_* values estimated for *S. norvegicus* for each combination of intervals of regions and depth.

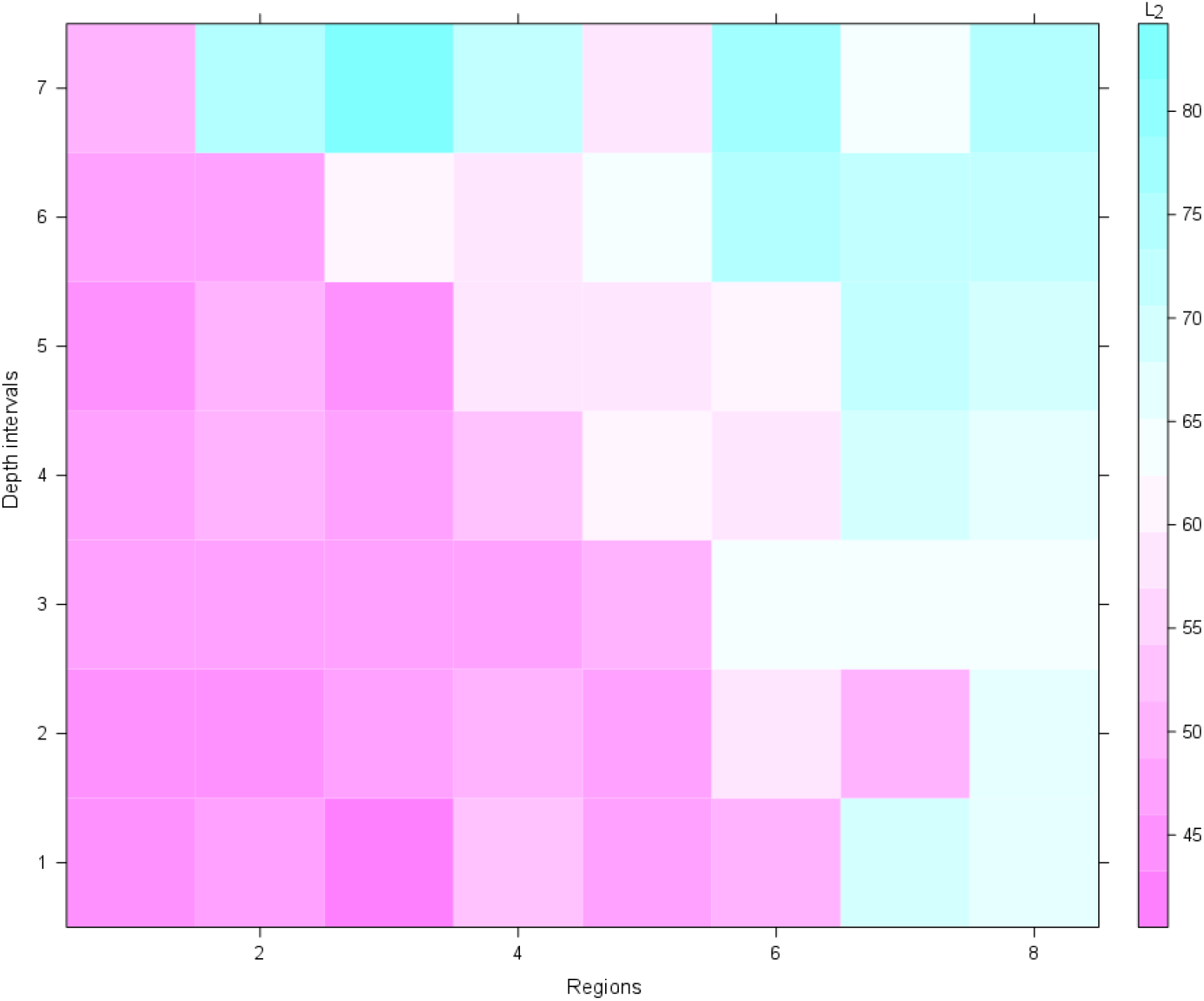

VBGF models with region as a main factor in the hierarchical model and depth as the second factor. Panels are for each combination of intervals of region/depth. Blue dots correspond to the observations. Blue line represents the model with only fixed effects common to the whole population. Pink line represents the mixed model effect with the region (fixed and random effects). Green line represents the hierarchical mixed model effect with regions as the first level and depth as the second one (fixed and random effects).

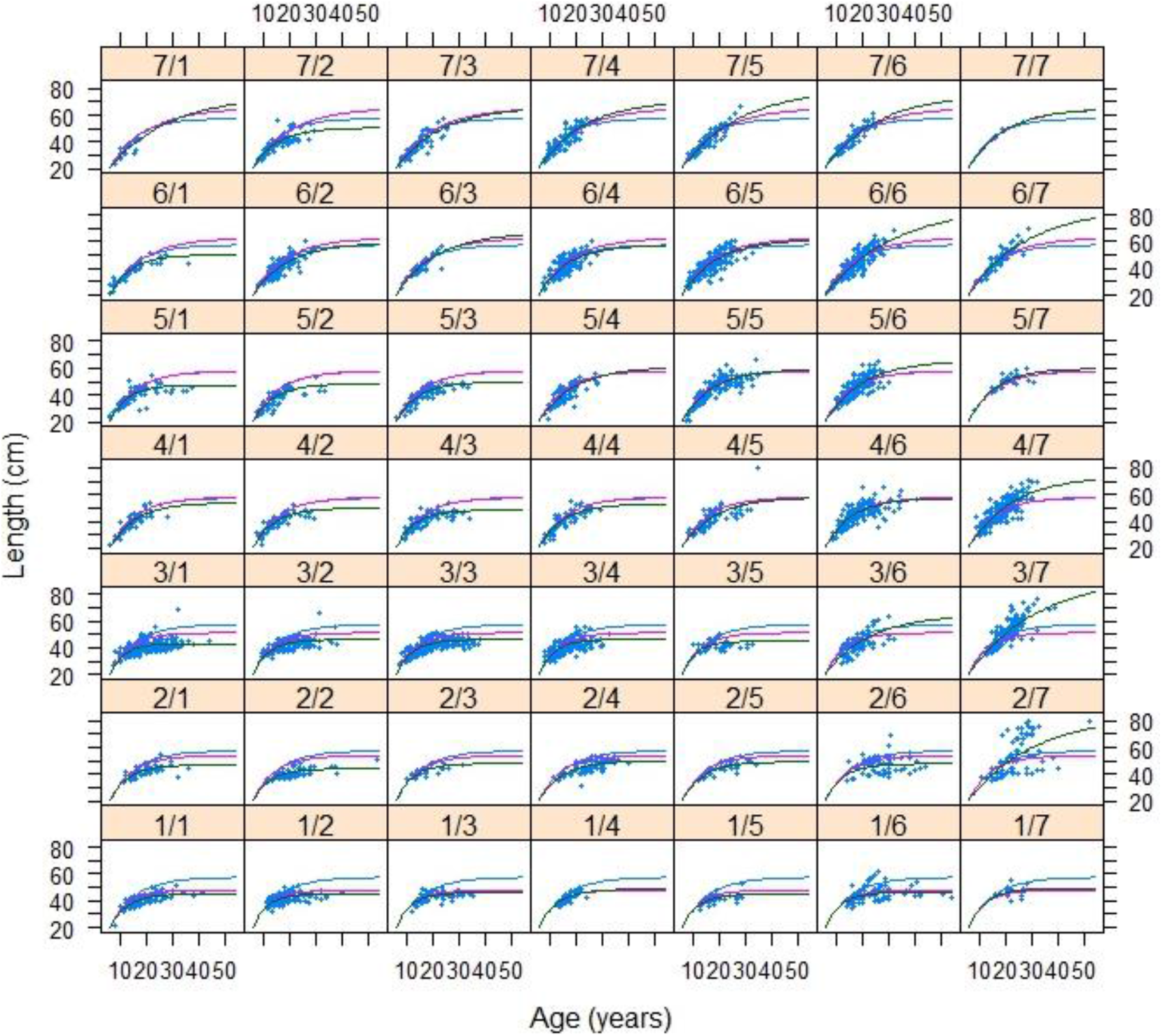

